# The effect of recognition on competition among clones in spatially structured microbial communities

**DOI:** 10.1101/2022.12.08.519541

**Authors:** Adrienna Bingham, Aparajita Sur, Leah B. Shaw, Helen A. Murphy

## Abstract

In spatially structured microbial communities, clonal growth of stationary cells passively generates clusters of related cells. This can lead to stable cooperation without the need for recognition mechanisms. However, recent research suggests that some biofilm-forming microbes may have mechanisms of kin recognition. To explore this unexpected observation, we studied the effects of different types of cooperation in a microbial colony using spatially explicit, agent-based simulations of two interacting strains. We found scenarios that favor a form of kin recognition in spatially structured microbial communities. In the presence of a “cheater” strain, strains with greenbeard cooperation was able to increase in frequency more than obligate cooperators. This effect was most noticeable in high density colonies and when the cooperators were not as abundant as the cheaters. We also studied whether a polychromatic greenbeard, in which cells only cooperate with their own type, could provide a numerical benefit beyond a simple, binary greenbeard. We found the greatest benefit to a polychromatic greenbeard when cooperation is highly effective. These results suggest that in some ecological scenarios, recognition mechanisms may be beneficial even in spatially structured communities.

## Introduction

Microbial life is rife with complex interactions between and within species. As a necessary part of existence, microbial social interactions can vary from chemical warfare, to competition, to synchronization, to cooperation (Tarnita, 2017; West et al., 2007). Like all cooperative communities, microbes are susceptible to invasion by selfish individuals who benefit from cooperation (Ghoul et al., 2014), but do not contribute (Velicer et al., 2000; Strassmann et al., 2000; Diggle et al., 2007; Sandoz et al., 2007; Zhang et al., 2009; Vos and Velicer, 2009; Popat et al., 2012). Despite the potential for invasion, cooperative behaviors are prevalent in natural systems. Such behaviors can often be explained through kin selection: individuals cooperate with relatives that likely share genes for the behavior. If the fitness benefits of the behavior are large enough, the frequency of the cooperative allele(s) will increase in the population, such that cooperation will outcompete other strategies (W. D. Hamilton, 1964a; W.D. Hamilton, 1964b).

One way to ensure the stability of kin-selected cooperation is through kin recognition in which individuals recognize and preferentially cooperate with related individuals (Waldman, 1988). While recognition of relatives has been documented in animals, it is uncommon in microbial systems. Instead, there is “kind” recognition where cooperators recognize each other through a particular signal; this system is known as a greenbeard. A greenbeard locus encodes a cooperative behavior (a single or multiple, tightly linked genes), a signal of cooperation, and the ability to recognize the signal in others (Gardner and West, 2010; Queller, 2011). For a greenbeard system to function, there must be relatedness at the locus, but not necessarily across the entire genome. Microbial examples include cell adhesion proteins that allow cells to attach to one another, as has been reported in yeast and social amoebae (Queller et al., 2003; Smukalla et al., 2008; Strassmann et al., 2011; Belpaire et al., 2022).

In aggregative cooperative behaviors that require motile microbes to locate one another before adhering, such as the formation of foraging slugs, fruiting bodies, or swarms, rare instances of kin- and self-recognition have been reported (Cao and Wall, 2017; Hirose et al., 2011; Stefanic et al., 2015; Vos and Velicer, 2009). Rather than a binary system of cooperation or non-cooperation, as is expected for a greenbeard locus, the proteins responsible for cellular adhesion exhibit a spectrum of cooperation (e.g., strength of cell-to-cell interaction) related to allelic variation, thus creating a “polychromatic greenbeard” (Smith, 1976; Gruenheit et al., 2017). A polychromatic greenbeard is more similar to traditional kin recognition, as individuals with identical “shades” of green tend to be more closely related across the genome, as long as rates of recombination are low (Jansen and van Baalen, 2006; Rousset and Roze, 2007).

In contrast to motile microbes, many microbes live and interact in stationary, spatially-structured communities known as biofilms. These communities are characterized by attachment to a surface, an extracellular matrix produced by the microbes, cellular differentiation, and increased resistance to environmental stressors (Costerton et al., 1995). Biofilms exist in nearly every environment in which microbes can be found, including medical settings where their presence is a particular risk due to their increased resistance to antimicrobials (Costerton et al., 1999; Hall-Stoodley et al., 2004). Biofilms require individuals to cooperate and produce goods that will be used by other members (i.e., extracellular matrix, drug efflux pumps, diffusible enzymes, quorum sensing molecules) (West et al., 2007). Mathematical modeling, simulations, and microbial experiments have demonstrated that unlike in aggregative and motile phenotypes, biofilms do not require a recognition system for cooperation to be stable (reviewed in (Nadell et al., 2016)). Clonal growth generates local patches of high-relatedness and leads to lineage sorting (Xavier and Foster, 2007; Tarnita et al., 2009; Nadell et al., 2010). In this way, spatial assortment passively leads to kin selection without the need for kin recognition. Despite the body of work suggesting recognition systems are not required for stable cooperation in biofilms, recent research has provided evidence that a cell adhesion protein may actually act in recognition in the budding yeast *Saccharomyces cerevisiae*, a stationary, biofilm-forming eukaryotic microbe (Oppler et al., 2019; Brückner et al., 2020).

Given this unexpected observation, here we ask whether greenbeard and polychromatic greenbeard systems could provide a benefit to cooperators in stationary microbial communities. While such recognition systems may not be required for stable cooperation, it is possible they provide enough of a fitness benefit to explain the presence of recognition systems in species whose characteristics preclude motility. We hypothesize that even with the passive emergence of clusters of direct kin, the ability to restrict cooperative benefits will lead to increased representation of cooperators in a growing spatially-structured community.

Our approach to addressing this question uses spatially-explicit, agent-based simulations of a growing microbial community containing two cell types. The simulations represent a simplified microbial community in which cells interact under different scenarios (with different sets of rules). Briefly, in our simulations, cooperators have a slower basal growth rate than non-cooperators, which represents the cost of producing non-diffusible/locally diffused public goods and other traits associated with biofilm formation. Cells that are adjacent to cooperators have an increased growth rate due to the benefit provided by the cooperative goods. A greenbeard recognition system is implemented by allowing cooperator cells to only increase the growth rate of cooperator neighbors. Finally, we investigate the effect of a polychromatic greenbeard by simulating microbial communities with two cooperator cell types that have different basal growth rates and can restrict growth benefit to their own cell type. We simulate different scenarios over a range of starting ratios of each cell type and of the strength of cooperation.

## Methods

We used an agent-based model to study the interactions between two cell types in a microbial community. The biological details were inspired by the yeast *Saccharomyces cerevisiae*, in which the potential for a recognition system has been reported (Oppler et al., 2019; Brückner et al., 2020). We used a published framework (Momeni et al., 2013), which we adjusted to apply to our research questions. The published model considered the effect two different cell types would have on one another, but did not consider the effect each cell type would have on other cells of its own kind. Our model is summarized in this section and the details of the simulation are given in the Supplementary Material. Briefly, to begin the simulation, cells were placed on a three-dimensional cubic array, and then stochastic Monte Carlo simulation was used to model their division. Social interactions were implemented by altering the rate at which a cell divided based on the occupancy and composition of the immediate neighborhood. When a cell divided, a daughter cell was placed at a nearby location. At the end of the simulation, the number and placement of each cell type was used for calculation.

Each cell type started with a baseline growth rate, *r*_*S0*_ and *r*_*F0*_. We assumed *r*_*S0*_ < *r*_*F0*_, or that there is a slower (*S*) and faster (*F*) growing cell type. For example, when only one cell type was capable of cooperation, we designated the cooperators *S* and the non-cooperators *F* to account for the costs of cooperation. In the presence of social interactions, the growth rates for the two cell types were:

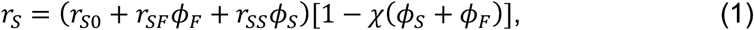

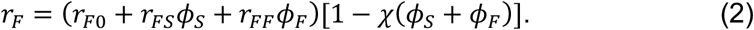

Parameters and variables are listed in Table 1 and are explained below.

| Parameters | Meaning/Purpose | Value |
| --- | --- | --- |
| $r_S$ | Growth rate of slow cell | Calculated in simulation |
| $r_F$ | Growth rate of fast cell | Calculated in simulation |
| $r_{S0}$ | Baseline growth rate of slow cell | 0.05 |
| $r_{F0}$ | Baseline growth rate of fast cell | 0.1 |
| $r_{SS}$ | Effect of slow cells on other slow cells | 0, 0.1, 0.3, 0.5, 0.7 |
| $r_{FS}$ | Effect of slow cells on fast cells | 0, 0.1, 0.3, 0.5, 0.7 |
| $r_{FF}$ | Effect of fast cells on other fast cells | 0, 0.1, 0.3, 0.5, 0.7 |
| $r_{SF}$ | Effect of fast cells on slow cells | 0, 0.1, 0.3, 0.5, 0.7 |
| $\chi$ | Competition effect | 1 |
| $\phi_S$ | Occupancy fraction of slow cells in neighborhood | Calculated in simulation |
| $\phi_F$ | Occupancy fraction of fast cells in neighborhood | Calculated in simulation |
| $N_c$ | Maximum biofilm length and width | 50 |
| $N_z$ | Maximum biofilm height | 100 |
| $T$ | Maximum number of simulated cells | 50,000 |
| $d$ | Length of inoculation square | 15 |
| $f$ | Fraction of inoculation square filled with cells | 0.05 |
| $f_S$ | Fraction of slow cells in inoculum | 0.1, 0.3, 0.5, 0.7, 0.9 |
| $R_i$ | Interaction radius | 3 |
| $R_d$ | Displacement radius | 3 |
| $\Delta t$ | Time step | 0.1 |

To determine the growth rate of an individual cell, we first calculated the local density of each cell type (fraction of sites occupied), *ϕ*_*S*_ and *ϕ*_*F*_. These were calculated for a cubic neighborhood within an interaction radius *R*_*i*_ surrounding the growing cell. A competition effect, *χ*, was incorporated and set to 1, such that the final term in the growth rate calculation, 1 – *χ* (*ϕ*_*S*_ + *ϕ*_*F*_), resulted in a zero growth rate when the neighborhood was full. Social interactions other than competition were set by the parameters *r*_*ij*_, where each of *i* and *j* was *S* or *F*. The parameter *r*_*ij*_ determined the effect that cell type *j* had on type *i*. For example, when strain *F* was a non-cooperator, its presence would have no effect on the slow strain or itself, so *r*_*FF*_ = *r*_*SF*_ = 0. However, when a strain was a cooperator, it would have a beneficial effect on others of its type, so *r*_*SS*_ > 0.

The growth rate parameters, *r*_*ij*_, were varied to simulate different social interactions and included some values that allowed the slower cells to achieve the same or greater growth than the non-cooperative fast cells. We considered four scenarios in our simulations: baseline competition, obligate cooperation, greenbeard cooperation, and polychromatic greenbeard cooperation. First, for baseline competition, all social interactions other than competition were absent (*r*_*SS*_ = *r*_*FF*_ = *r*_*SF*_ = *r*_*FS*_ = 0). Next, for obligate cooperation, both *r*_*SS*_,*r*_*FS*_ > 0, while *r*_*FF*_,*r*_*SF*_ = 0. We set *r*_*FS*_ = *r*_*SS*_ so that non-cooperators derived the same benefit from cooperator neighbors as a cooperator would. This was the simple case in which non-cooperative cheaters could benefit from the presence of cooperators. In the third scenario, a greenbeard cooperator, we again set *r*_*SS*_ > 0.

However, the cooperator could now restrict its benefit to its own cell type, and the non-cooperator strain did not benefit from the public goods, so *r*_*FS*_,*r*_*FF*_,*r*_*SF*_ = 0. Finally, in the polychromatic greenbeard scenario, both the fast and slow strain were cooperators that were able to restrict their cooperation to their own strain, *r*_*SS*_ = *r*_*FF*_ > 0, and *r*_*SF*_,*r*_*FS*_ = 0. To determine if a polychromatic greenbeard was beneficial, for comparison, we also considered the situation where both the fast and slow cooperators were simple greenbeards that do not restrict cooperation, *r*_*SS*_ = *r*_*FF*_ *= r*_*SF*_ = *r*_*FS*_ > 0.

The simulations began with a diluted inoculum of cells into a small square of size *d* × *d* at the center of the bottom layer of our cubic array (see Fig. 1A), similar to a migration event in the environment or a droplet on a petri dish. This allowed us to simulate outward expansion of an initial community/colony from a founding event, in contrast to the simulation in Momeni et al. (2013) with random inoculation throughout their whole bottom layer. A fraction *f* = 0.05 of the sites were randomly selected to fill with cells, and a fraction *f*_*S*_ of the initial cells were assigned as the cell type with slow baseline growth, with the remainder as the faster type.

**Figure 1:**
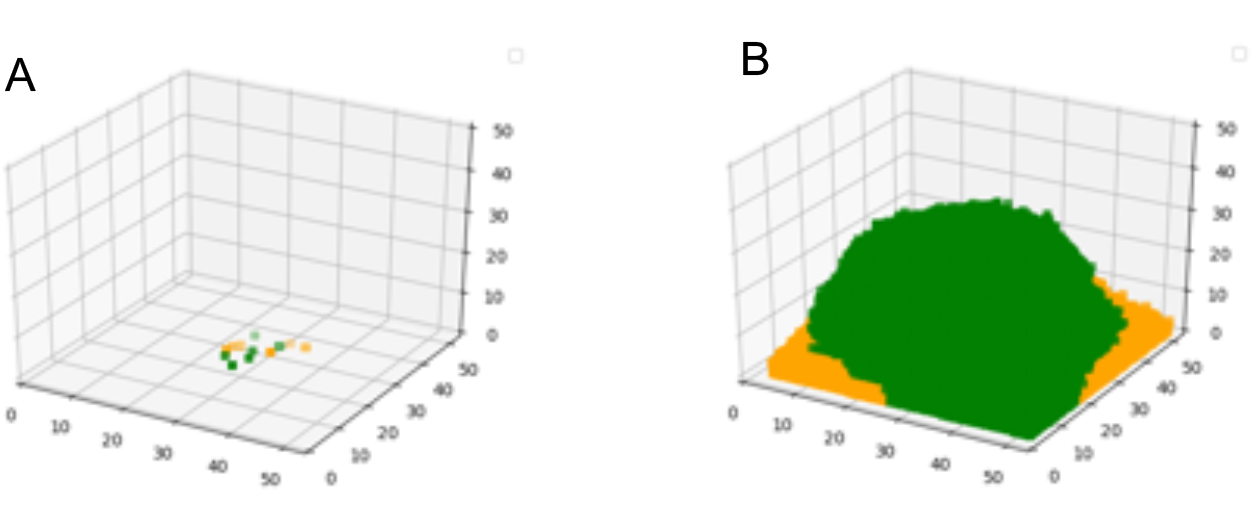
(A) Schematic of cells randomly distributed within a centered square droplet on the bottom surface of the cubic array in the simulation. The green squares are slow/cooperative cells and the orange squares are fast/non-cooperative cells. This figure depicts a cell inoculation with *f*_*S*_ = 0.5, or an equal proportion of each cell type. (B) Schematic of cells at the end of a simulation for which *f*_*S*_ = 0.5, *r*_*SS*_ = 0.5, and *r*_*FS*_,*r*_*FF*_,*r*_*SF*_ = 0, a greenbeard cooperator scenario.

When a cell divided, the next step was to find an empty site for the daughter cell to be placed. If there was an empty site within a square neighborhood of radius *R*_*d*_ in the same horizontal plane, the cell divided horizontally, pushing other cells aside as needed (see Supplementary Material, Figures S1-2). A similar mechanism to push nearby cells was implemented by Momeni et al. (2013). While we set the value of *R*_*d*_ to 3, their data suggested that the value could be as high as 5. If there were no empty sites in the displacement neighborhood in the same horizontal plane, the daughter cell was placed directly above the mother cell as this has been shown to be a characteristic of *S. cerevisiae* (Momeni et al., 2013). If there were cells above the mother cell, they were pushed upwards.

Cell growth was simulated until a specified stopping point was reached. In most cases, the stopping point was when the community reached a total of 50,000 cells (see Figure 1B), which, through multiple simulations, we determined represented a full community. At this point, the bottom of the grid was over 90% filled and cells throughout the community had begun to divide upwards, generating a three-dimensional structure.

The total number and location of each cell type were stored when a cell in the bottom layer first touched the outer edge. This was intended to capture the community during growth when population structure was forming, but the community was still sparse. Data were also stored at the end of the simulation. In addition to storing the total proportion of each cell type, we stored the proportion of each type at the surface of the colony. To define the colony surface, we began at each point in the top layer of the cubic array and counted downward until we reached the topmost cell. We ran 20 simulations for each combination of parameters; measurements were averaged over all simulations.

As noted above, the growth rate depended on both the social interaction parameters and the neighborhood surrounding the cell. Having a small number of cooperator neighbors increased growth but having too many could reduce growth due to competition. The advantage of increasing the cooperation parameter *r*_*SS*_ was thus not directly obvious. By differentiating Eq. 1 with respect to occupancy fraction, we found that the maximum growth rate a slow strain could achieve, attained when 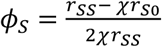 and *ϕ* = 0, is 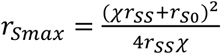. In our simulations, we set *χ* = 1. For the parameter values used here, the maximum slow cell growth rate *r*_*Smax*_ was monotonically increasing with the cooperation parameter *r*_*SS*_. For easier interpretation of our results, we plotted our simulation results versus the maximum possible growth rate of slow cells rather than the bare parameter *r*_*SS*_. Note that the maximum growth rate for baseline competition was 0.05, or the basal growth rate for slow cells. For all other simulations in which the slow cells were cooperators, the maximum growth rate exceeded the basal growth rate.

## Results

### Cooperation in a Spatially Structured Community

To investigate the dynamics of the different social scenarios, we began by looking at the proportion of slow cells at the end of the simulations (Figure 2). When compared to the initial proportion in the inoculum, the final proportion can reveal an increase or decrease in frequency of the slow strain throughout growth of the spatially structured community. We also considered the proportion of each cell type on the surface of the community, as dominating the outer edge of an expanding front provides increased access to nutrients and resources, and can be more important for fitness than overall presence throughout the community (Hallatschek et al., 2007). We found that proportion of cells on the surface closely mirrored total proportion (Figure 2A), so we present only the total proportion in the rest of the analyses.

In the baseline scenario with no cooperation, and only a fast and a slow growing strain, the clear expectation is that the slow growing strain will be outcompeted by the end of the simulation. This can be seen in the first data point (black square) in each panel in Figure 2A, when *r*_*Smax*_= 0.05, which is simply the baseline growth of the slow strain (because *r*_*SS*_ = 0). In all cases, regardless of the initial proportion of slow cells in the inoculum, the slow strain was nearly non-existent by the end of the simulation.

When cooperation was introduced by allowing slow cells to provide a benefit to other slow cells (*r*_*SS*_ > 0), as expected, the slow strain remained in the community throughout the simulation and, in some cases, even increased in frequency. The results of obligate cooperation (when slow cells/cooperators provide a benefit to all nearby cells) can be seen in purple in Figure 2. Note that in this scenario, cooperator cells also increased the growth rate of non-cooperator cells, allowing “cheating”. When cooperation was very strong (*r*_*Smax*_ = 0.15 or 0.20), the cooperator genotype was able to increase in frequency compared to its initial frequency. Thus, these results qualitatively support the previous work that showed that clonal growth in a spatially structured community can favor cooperation, even when cheating is possible, as long as public goods are only locally diffused (Nadell et al., 2010; Xavier and Foster, 2007).

**Figure 2A:**
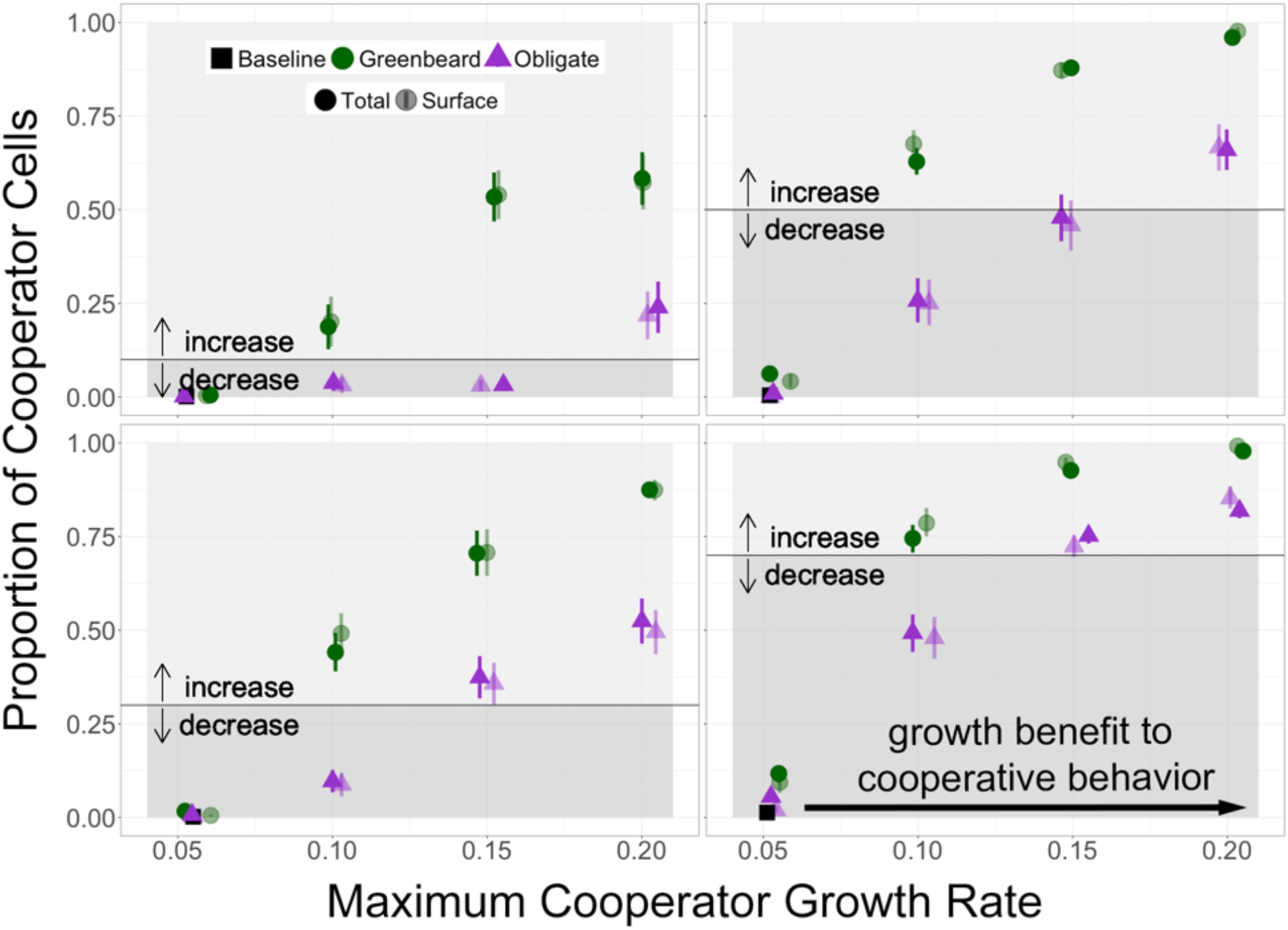
Proportion of the slow, cooperator strain at the end of simulations in the total colony and on the outer surface of the colony over a range of cooperative parameter values (r_SS_ = 0.1, 0.3, 0.5, 0.7). Purple points represent obligate cooperation, where the slow strain provides a benefit to all nearby cells. Green points represent greenbeard cooperation, where the slow strain provides a benefit only to other slow cells. Solid points represent the total final proportion of cooperators; transparent points represent the final proportion of cooperators at the top of the community. Baseline growth rate is the first data point, *r*_*Smax*_ = 0.05. Error bars are standard error of the mean. Panels represent a range of initial proportions *f*_*S*_ of the slow strain, indicated by the black lines; arrows represent an increase or decrease of the cooperator strain compared to the initial frequency. Data points are jittered for clarity and are plotted only for the four specified r_SS_ values.

In the next set of simulations, the slow strain was again a cooperator, but this time was able to restrict its benefit to other cooperators, thus simulating a greenbeard (green points in Figure 2). In this scenario, when the maximum possible growth rate of the cooperator was equal to the baseline growth rate of the non-cooperator (*r*_*F*0_ = 0.1), the final proportion of cooperators was greater than its initial proportion. This suggests a benefit to a greenbeard recognition system, even in a spatially structured community. Compared to the obligate cooperator scenario, as the initial fraction of slow cells increased (comparing the four panels), the final proportion of slow cells saturated more quickly with increasing maximum growth rate. Overall, the frequency of slow cells was higher when expressing greenbeard cooperation than when non-cooperators were able to benefit from their public goods. When visualizing the resulting communities in the two types of simulations (Figure 2B), a difference in spatial distribution is apparent.Communities with obligate cooperators appear to be more mixed than communities with greenbeard cooperators, which appear to have cooperators spatially separated. This makes intuitive sense, as obligate cooperators are providing benefits to non-cooperators, while greenbeard cooperators are excluding non-cooperators. When a greenbeard is viewed as the ability of cells to adhere to one another, this spatial exclusion supports the observation that adherence can be an advantageous trait in microbial competition (Schluter et al., 2015).

**Figure 2B:**
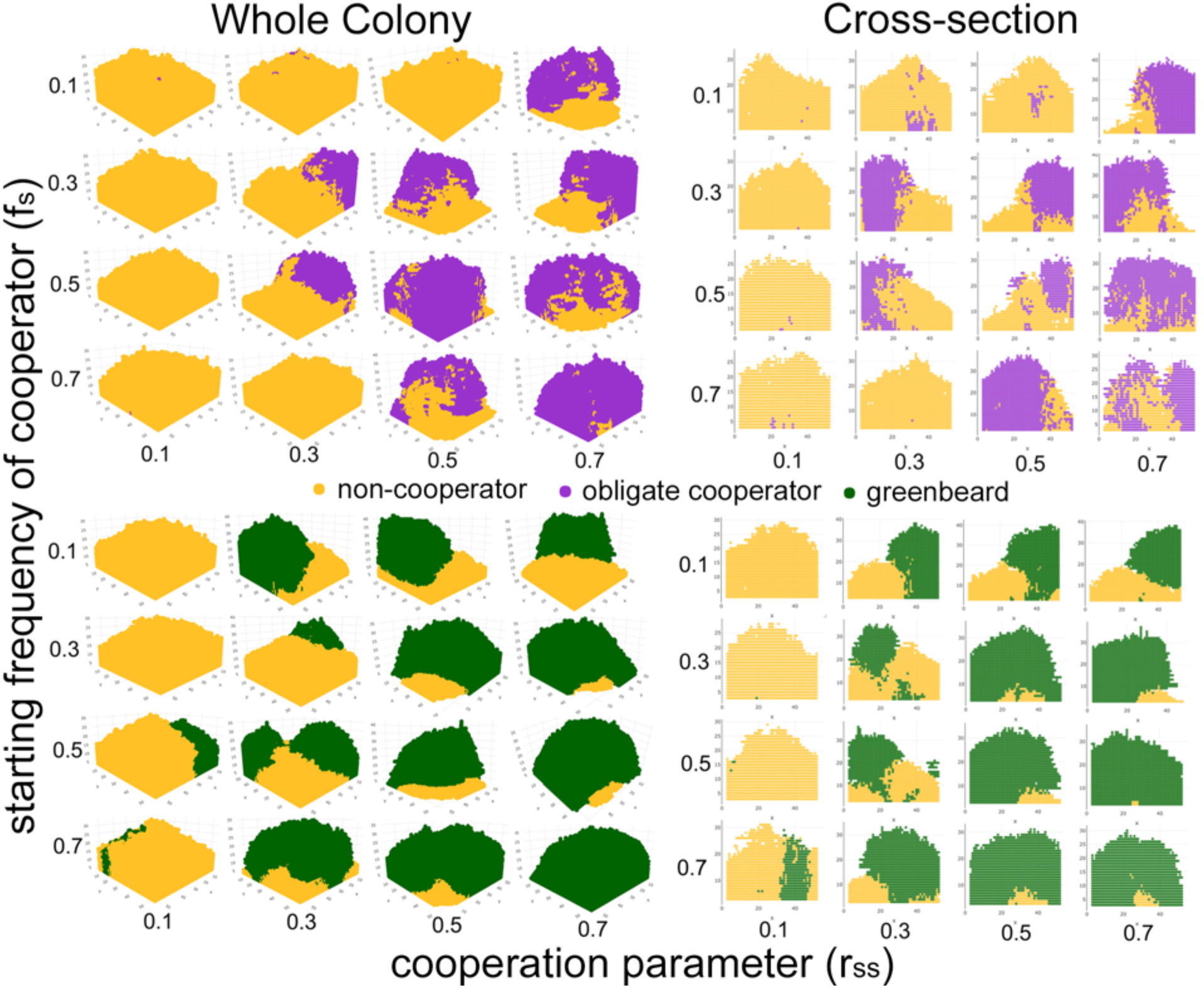
Sample simulated colonies (left) and cross-sections (right) over a range of cooperative parameter values and cooperator starting frequencies. The first simulated colony for each combination of parameters was plotted in the 3-dimensions used by the simulation (x, y, and z axes), as well as a 2-dimensional vertical slice from the middle of the colony (at y = 25). Yellow points represent non-cooperators, purple points represent obligate cooperators, and green points represent greenbeard cooperators.

### A Comparison of Obligate and Greenbeard Cooperation

To determine how much the slow strain benefited from restricting its cooperation (i.e., expressing a greenbeard recognition signal), we looked at the difference between the total final proportion of slow cells in greenbeard and obligate cooperation (Figure 3), i.e., subtracting the purple points in Figure 2 from the corresponding green points. The results suggest that cooperator cells gained more of a benefit from a greenbeard system when they were a moderate fraction (roughly 50%) of the cells present. When there were more cooperator cells (i.e., when *f*_*S*_ = 0.7), there were not as many non-cooperators to take advantage of slow cells, so restricting cooperation had less of an effect. On the other hand, when the cooperators were initially rare, there was a larger benefit to restricting cooperation as long as cooperation was strong enough. Especially when the initial proportion of slow cells was 10%, the benefit to restricting cooperation was low for smaller values of max growth rates.

**Figure 3.**
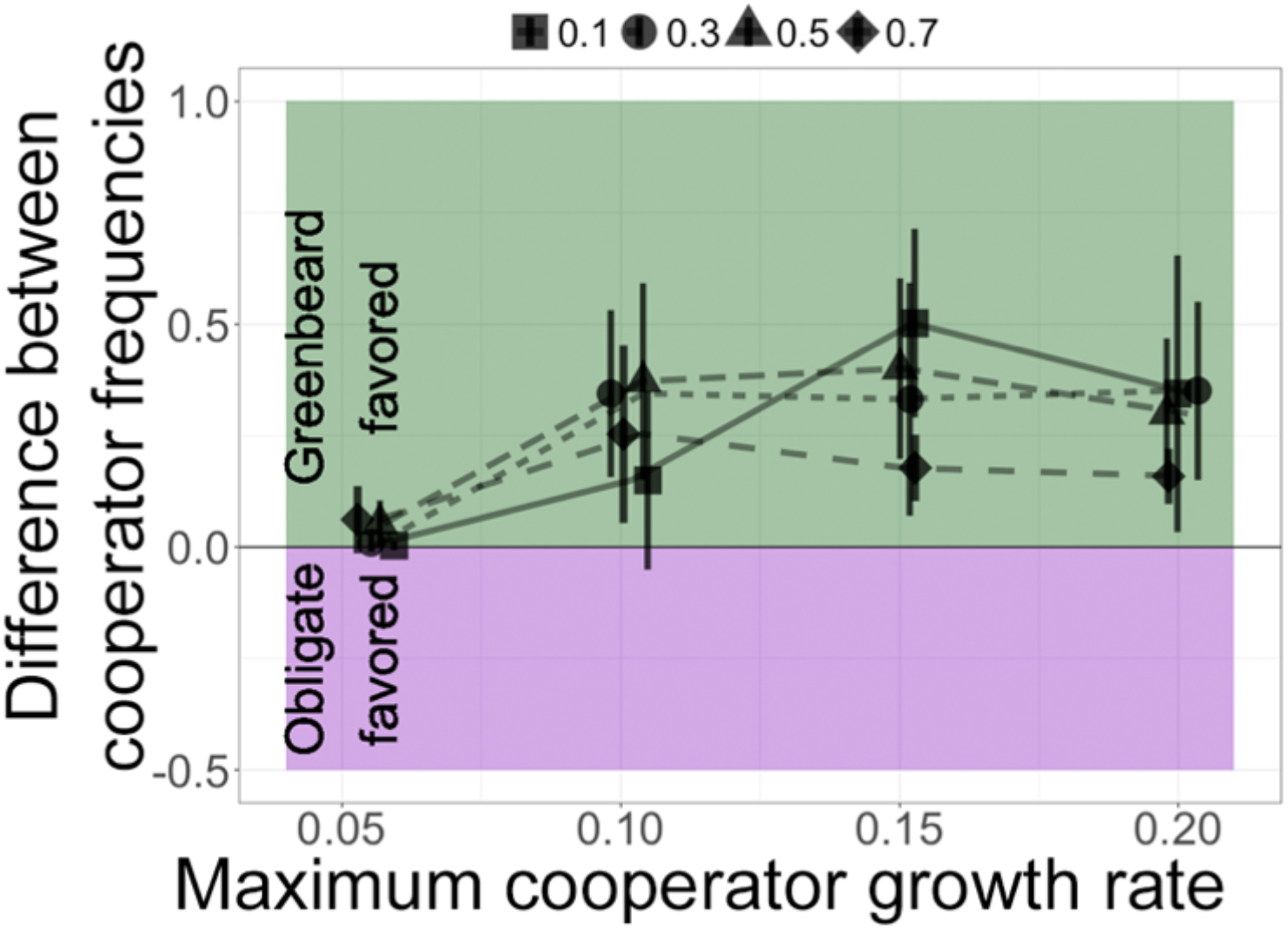
Comparison of types of cooperation when in competition with non-cooperators for a range of initial proportions and maximum cooperator cell growth rates. A value of 0 indicates the same final frequency of cooperator cells in obligate and greenbeard cooperation scenarios. Values above 0 indicate a higher final frequency of cooperator cells in the greenbeard scenario; error bars are standard error of the mean. Initial cooperator frequencies: 0.1-square and solid, 0.3-circles and dotted, 0.5-triangles and dashes, 0.7-diamonds and long dashes. Data points are jittered for clarity.

### Cooperation in Sparse Versus Dense Communities

Next, we investigated how the density of the communities affected the benefit of the two different types of cooperation. As shown in the previous two sections, when the communities were fully grown, there was an apparent benefit to obligate cooperation and a further benefit to greenbeard cooperation. This may be in part because the community has grown to a dense enough state for cells to have fuller neighborhoods and the opportunity to interact with one another. To test whether or not the density of the community had an effect on the benefit to cooperation, we compared the proportion of slow cells when the community was sparse and growing exponentially (i.e., when the first cell reached the outer edge of the cubic growth area) with the proportion of slow cells in a dense community at the end of the simulation (i.e., when the community had reached 50,000 cells).

As predicted, in general, slow cells had a higher total proportion at the end of the simulations than in sparse colonies (Figure S3). This suggests that the growth benefit to both obligate and greenbeard cooperation appear later in the simulation and in more dense conditions. This is likely because in a denser community and after a period of initial growth, slower cooperative cells are surrounded by more of their own kind and are able to benefit from each other.

We compared the benefits of obligate and greenbeard cooperation in sparse and dense conditions (Figure 4) and found that for all starting frequencies of cooperators, the benefit to greenbeard cooperation was generally greater in dense than sparse colonies. This is due to a higher chance of slow cells being near fast cells in denser colonies, and therefore a greater benefit to restricting cooperation and excluding fast cells. We also expect the greatest benefit to greenbeard cooperation when each cell type is present in equal numbers, since there are likely to be more interactions between the types. Indeed, conditions where the final proportion of each cell type is on the order of 50% (see Figure 2B) are associated with the highest benefit to greenbeard cooperation (Figure 4).

**Figure 4.**
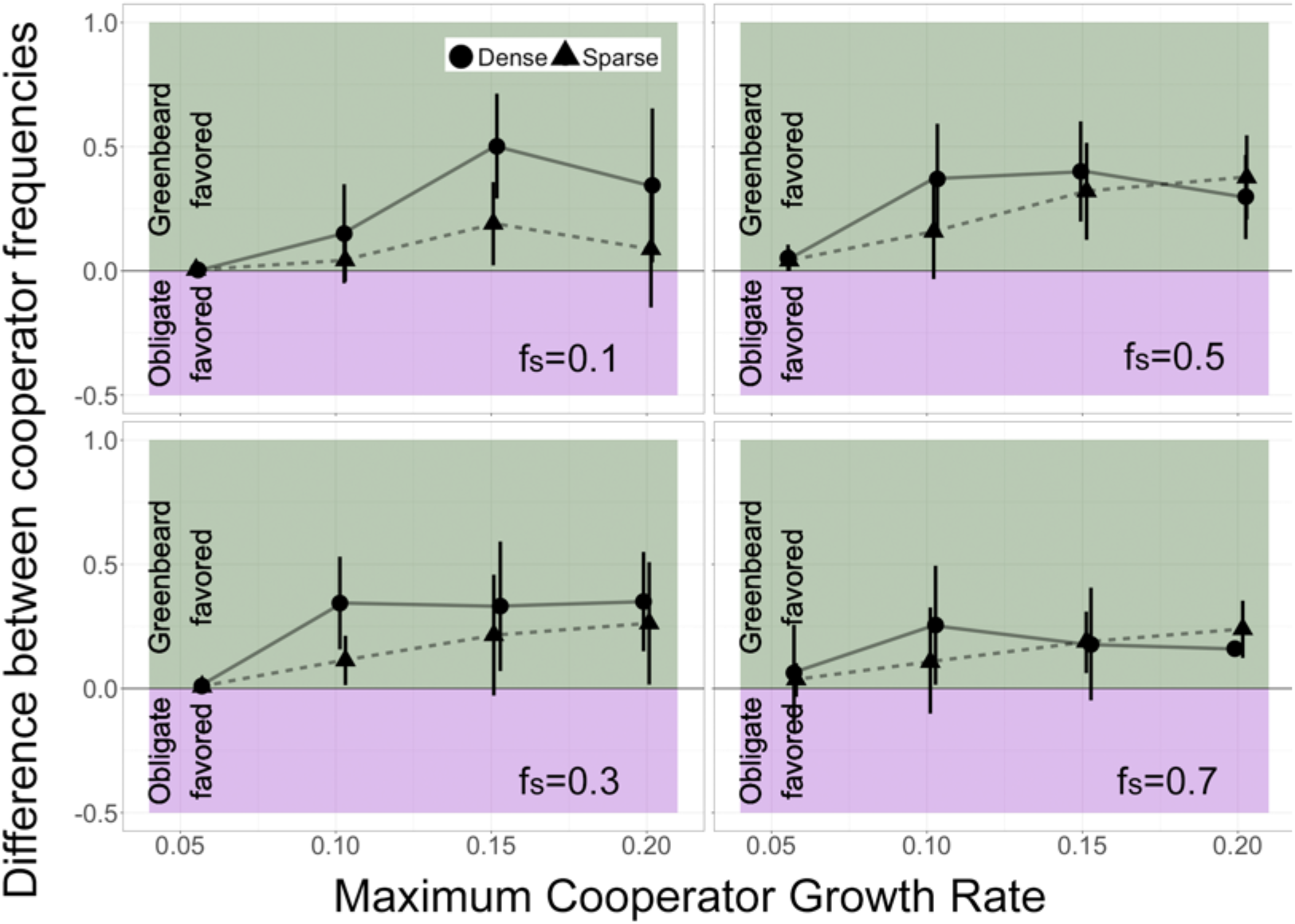
Benefit to greenbeard cooperation when colonies are sparse (dashed) or dense (solid). Comparison of types of cooperation when in competition with non-cooperators in sparse (triangles and dashed lines) and dense (circles and solid lines) growing conditions for a range of initial proportions and maximum cooperator cell growth rates. Value interpretation as in Figure 3.

### A Comparison of Greenbeard and Polychromatic Greenbeard Cooperation

The first set of simulations and analyses showed that greenbeard cooperation could increase in frequency more than obligate cooperation under certain conditions in spatially structured communities. We next sought to determine whether a polychromatic greenbeard (i.e., “true” kin recognition, or recognition of close relatives) could provide a further benefit over that which is provided by a general greenbeard. In natural circumstances, microbial lineages of the same species can have different baseline growth rates due to genetic differences. Even if a community contains only cooperating genotypes, it may benefit faster growing lineages to restrict cooperation in order to not be hampered by the slower growing lineages. This might explain why such a recognition system is observed in real species. To test this hypothesis, we again simulated communities with a fast and a slow growing strain, but this time, both were cooperative and we varied combinations of obligate and restricted cooperation (i.e., simple greenbeard and polychromatic greenbeard).

Figure 5 presents the results of simulations of four scenarios: (1) a fast and slow greenbeard cooperator, (2) a fast greenbeard and a slow polychromatic greenbeard, (3) a fast polychromatic greenbeard with a slow greenbeard, and (4) a fast and slow polychromatic greenbeard cooperator. The scenarios are summarized in Figure 6A. In all cases, we expected the fast strain to ultimately outcompete the slow strain; however, we explored whether certain scenarios enhanced or thwarted this process. A further starting ratio was added, *f*_*S*_ = 0.9, in order to clearly see the effect of the social scenarios. In all cases, we set *r*_*FF*_ *= r*_*SS*_, so both strains produced the same amount or quality of public good.

**Figure 5A:**
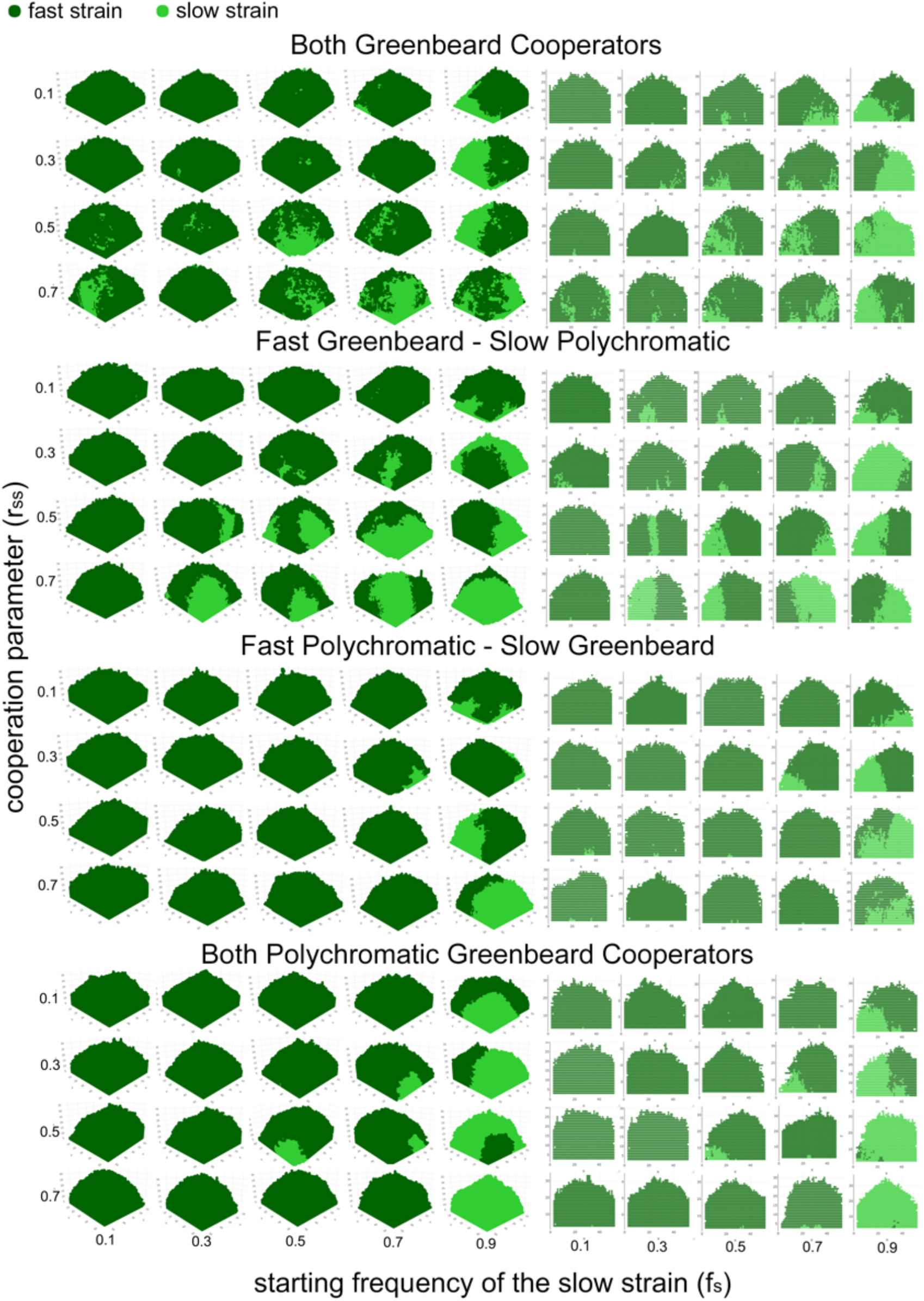
Sample simulated colonies (left) and cross-sections (right) over a range of cooperative parameter values and starting frequencies. The first simulated colony for each combination of parameters was plotted in the 3-dimensions used by the simulation (x, y, and z axes), as well as a 2-dimensional vertical slice from the middle of the colony (at y = 25). Dark green points represent the fast strain and light green points represent the slow strain.

**Figure 5B:**
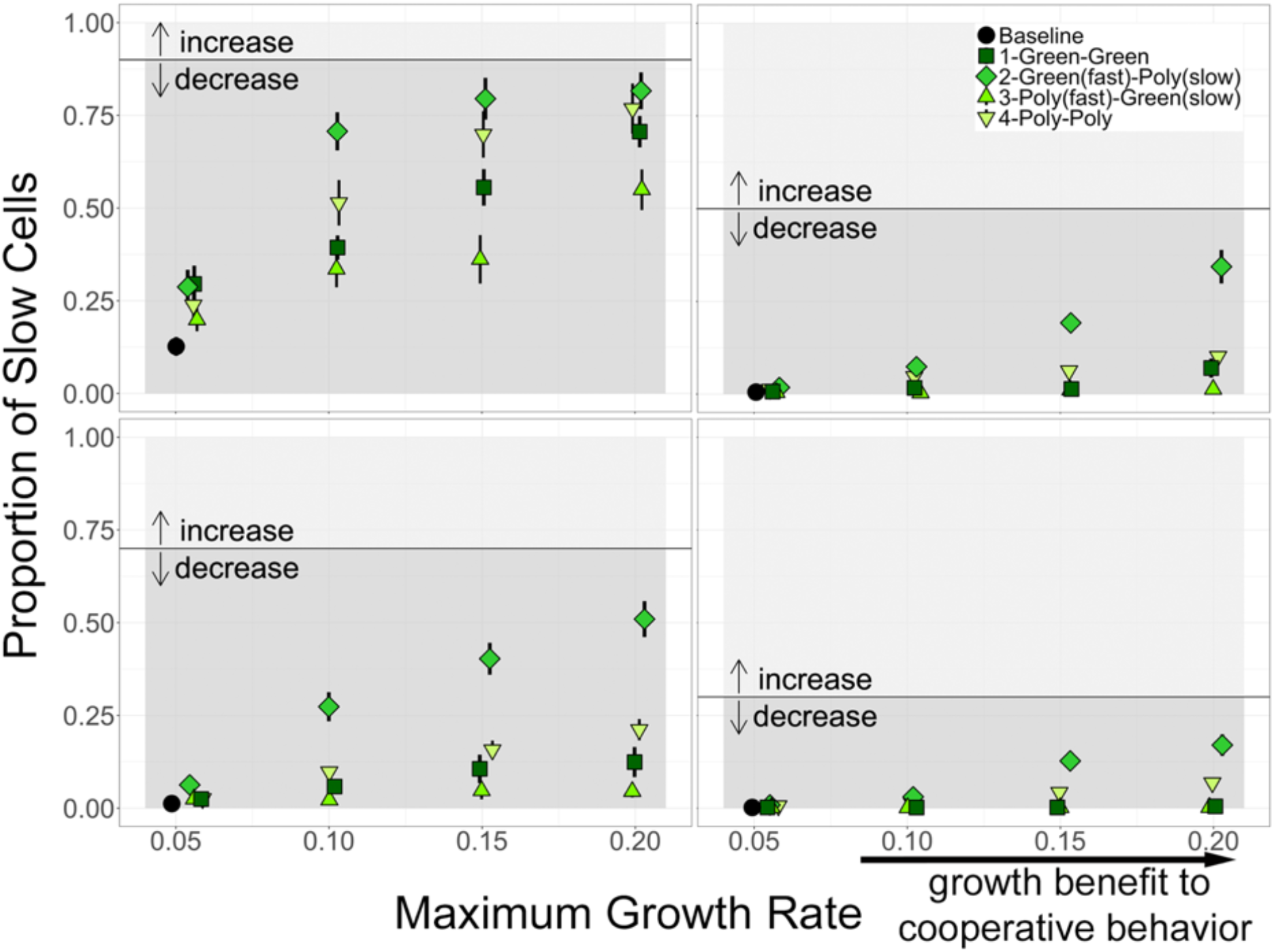
Total proportion of slow cells at the end of the simulations vs maximum growth rate of the slow strain. In each case, *r*_*FF*_ *= r*_*SS*_. For Scenario 1, both strains are simple greenbeards (i.e., they cooperate with both cell types). For Scenario 2, the fast strain is a simple greenbeard, but the slow strain is a polychromatic greenbeard (F cooperates with S, but S does not cooperate with F). Scenario 3 is the opposite, the fast strain is a polychromatic greenbeard, while the slow strain is simple greenbeard. For the last scenario, both strains are polychromatic greenbeard, so cooperate with their own, but not the other, cell type. The black lines are the initial proportion the focal strain. Error bars are standard error of the mean; points are jittered for visual clarity.

**Figure 6:**
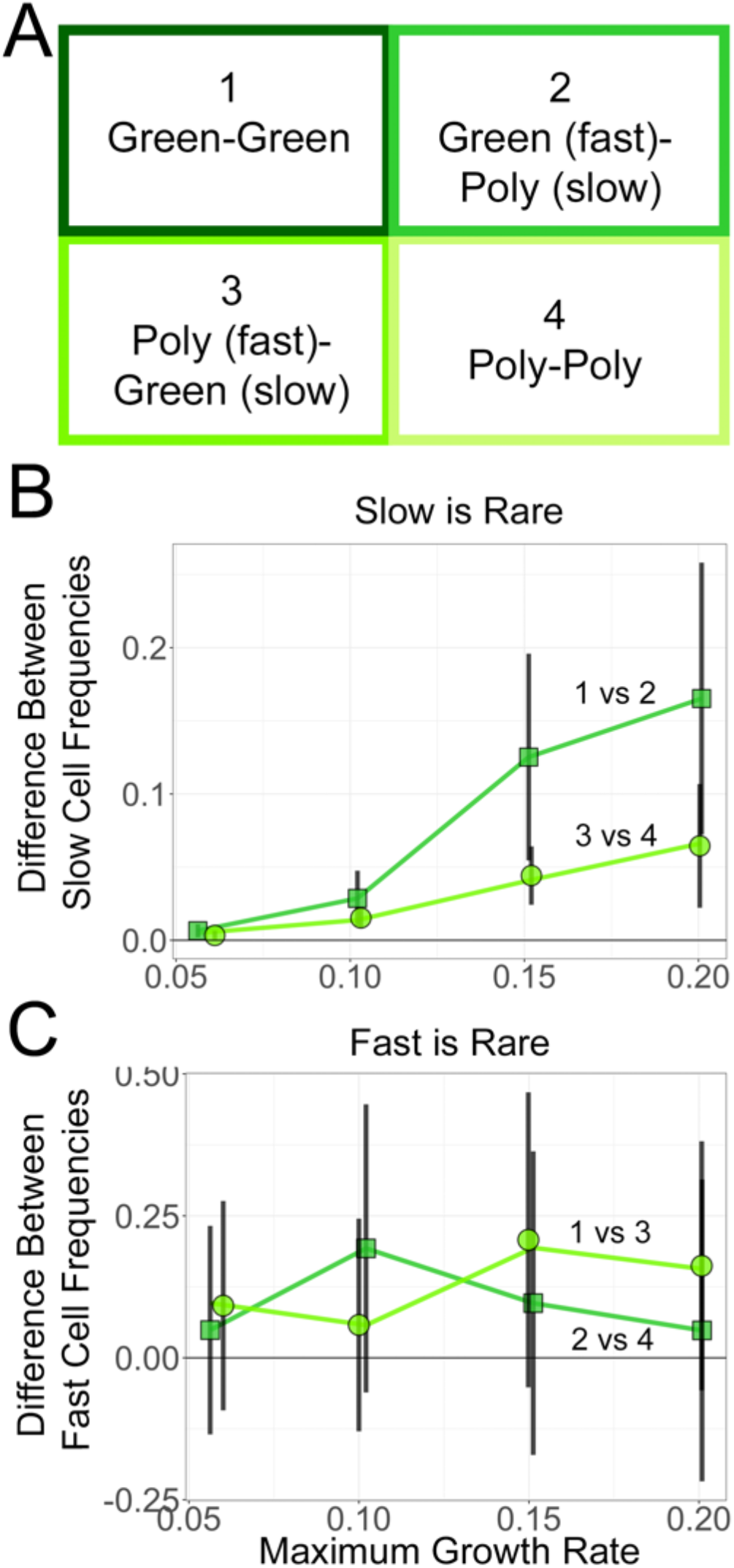
The effect of restrictive cooperation in cooperative communities. (A) Schematic of cooperative relationships between fast and slow cell types. (B) Comparison of final slow frequencies when slow cells gain restrictive cooperation in simulations begun at 0.3 slow cell frequency. (C) Comparison of final fast frequencies when fast cells gain restrictive cooperation in simulations begun at 0.1 fast cell frequency. Error bars are standard error of the mean; data points are jittered for clarity.

The baseline of no cooperation can again be seen in the first data point in each panel of Figure 5B (*r*_*imax*_ = *r*_*i0*_, or baseline growth), where the fast strain has outcompeted the slow strain. Looking at the four social scenarios that include combinations of cooperation, we see that the slow strain decreased in frequency over the timeframe of the simulations, regardless of the starting ratio: its long-term fate is presumably, and not surprisingly, to be outcompeted. However, the slower strain did remain a noticeable fraction in many of the simulations, which can be seen visually in the colony renderings in Figure 5A. As could have been anticipated, in all the scenarios investigated, the results are typically more pronounced the larger the cooperative benefit (i.e., as the maximum possible growth rate from cooperation increased).

We considered, in turn, the benefit to either the slow strain or the fast strain in developing the polychromatic trait, i.e., in restricting its cooperation. The slow strain became polychromatic as it transitioned from scenario 1 to 2 (when competing against a fast greenbeard) or from 3 to 4 (when competing against a fast polychromate). The benefit to the slow strain was greatest when the fast strain was a greenbeard and when cooperation was stronger. Most relevant is the case where the slow strain was initially rare (lower right panel of Figure 5B; Figure 6B), as one can observe whether a new, rare polychromatic mutation competed successfully. We saw that only when the fast strain was a greenbeard that provided substantial cooperation did the slow strain benefit much from becoming polychromatic. Otherwise, the slow strain was too heavily outcompeted by the fast strain.

The fast strain became polychromatic when it transitioned from scenario 1 to 3 (competing against a slow greenbeard) or from 2 to 4 (competing against a slow polychromate). The fast strain was initially rare in the upper left panel of Figure 5B and Figure 6C. Gaining a polychromatic trait was beneficial to the fast strain, regardless of whether the slow strain was a greenbeard or polychromatic, and especially when cooperation was stronger. From the subsequent panels in Figure 5B, we see that as the fast strain competed against a slow polychromatic strain, the benefit to the fast strain restricting its cooperation continued to be apparent. However, a slow greenbeard was a poor enough competitor that the fast strain gained less by restricting its cooperation.

In the colony renderings in Figure 5A, the type of cooperation led to some visual differences in the colonies. When both strains were greenbeards, the two strains appeared more intermixed. When both were polychromatic, each strain appeared more clustered with its own kind. However, the intermixing in the two-greenbeard case was probably less than in Momeni et al. double-cooperatorscenario because our greenbeards cooperated with their own cell type in addition to the other type, obviating the need for each cell to grow near cells of the other type. The effect of cooperation type on colony structure would be an interesting topic for further study.

## Discussion

The goal of the simulations presented here was to investigate the effects of different types of cooperation on the frequency and arrangement of cells in spatially structured microbial communities, and in particular, determine whether a polychromatic greenbeard (i.e., microbial kin recognition) could provide a fitness or growth benefit in such a community. Our simulations began with a small inoculum of different starting frequencies of two cell types. This could represent an experimental situation, such as starting a colony on a petri dish, or a more natural situation, such as a migration via an insect to a new environment or resource patch. The probability of growth for each cell depended on its type and neighborhood. We tracked the colony growth over time and monitored the change in frequency of the cell types. The simulations were fitness-based models that altered growth rates based on different types of cell-cell interactions. The model did not consider biological mechanisms, such as nutrient flux or enzyme diffusion, as the goal was to determine whether certain types of interactions could provide a fitness benefit, not to model the specific dynamics of an experimental system (Nadell et al., 2013, 2016).

Despite not including explicit mechanisms in our model, the scenarios we simulated are rooted in biological phenomena that exist in spatially structured communities. The obligate cooperators growing with non-cooperators captures the idea of a locally diffusible enzyme that can be used by producers and non-producers alike (e.g., Greig and Travisano, 2004; Griffin et al., 2004; Harrison, F and Buckling, A, 2009; Kümmerli, R et al., 2014; Lindsay et al., 2018), or floating aggregates that can contain non-producing cells that ultimately lead to the collapse of the community (Rainey and Rainey, 2003). It is within this context that it is possible to ask whether a recognition mechanism in which cooperators only provide growth benefits to other cooperators might provide a fitness advantage, which was tested in our greenbeard and non-cooperator simulations. For biofilms, these simulations capture the idea of only adhering to cells also expressing adhesins (Smukalla et al., 2008; Belpaire et al., 2022). And finally, we ask if restrictive cooperation in which cooperators provide growth benefits exclusively to specific cooperator types might provide a further fitness advantage, tested in our polychromatic greenbeard simulations. In the case of diffusible enzymes, this could be the advantage of enzyme-receptor specificity (Butaitė et al., 2017), while in the case of biofilms, it could be adhering to cells with the same adhesin alleles (Brückner et al., 2020).

In the first set of simulations, our results confirmed that obligate cooperation can increase in frequency in spatially structured communities, even when “cheating” (Ghoul et al., 2014; Travisano, M and Velicer, GJ, 2004) is possible, as long as the growth benefit provided by cooperative cells is only local. This has been studied most extensively as diffusible public goods using experimental, computational, and theoretical approaches (reviewed in Nadell et al., 2016). Our simulation model results qualitatively agree with the conclusions of those studies: when growth promotes clonal clusters, the benefit to the cooperative behavior is mostly shared with clonemates, and can therefore resolve the public goods cheating dilemma.

In this first set of simulations, we also investigated greenbeard cooperation in a community with non-cooperators. In microbes, this kind of cooperation is usually in the form of cell-cell adherence, and has been investigated experimentally and computationally (Xavier and Foster, 2007; Smukalla et al., 2008; Nadell et al., 2010; Schluter et al., 2015; van Gestel and Wagner, 2021; Belpaire et al., 2022). Once again, our simulations qualitatively confirm findings from these studies: the ability to adhere selectively promotes spatial segregation and can even be used as a weapon against non-adhering strains.

In our simulations, we were able to compare the increase in frequency of cooperators under the different types of cooperation, and found that greenbeard cooperation is able to increase in frequency more than obligate cooperation, a somewhat surprising result for spatially structured communities. The clonal growth of stationary cells passively generates local clusters, so it was unclear in advance whether a greenbeard cooperator would provide much of a growth advantage compared to an obligate cooperator that is able to interact within a cluster of like types. Finally, we found that the benefit to both obligate and greenbeard cooperation when grown with non-cooperators was realized as communities became denser at the end of the community expansion. The increase in density allowed the growth benefits of cooperators to have an effect as cells were in contact more (Yanni et al., 2019).

Our second set of simulations investigated whether a polychromatic greenbeard, in which the cells only cooperate with their own type, could provide a numerical benefit beyond a simple, binary greenbeard. The results showed that a more fit, faster-growing lineage will be hampered by the presence of a less fit, slower-growing lineage because the slower lineage will gain a growth benefit from being near the more abundant, faster one. The results also show that if one of the strains were able to restrict their cooperation (the asymmetric case of one polychromatic greenbeard and one simple greenbeard), such a strategy would be favored over the symmetric greenbeard case (i.e., the faster strain increases its relative frequency when it is selectively cooperative, and the slower strain increases its relative frequency when it is selectively cooperative). Finally, we also see that restoring the symmetry, by both strains only cooperating with their own type, would be favored. That is, when one strain has the advantage of restricting cooperation (i.e., a form of “cheating”), if other strain also becomes selectively cooperative, its frequency will increase. This suggests that if one lineage were to evolve a mechanism for selective recognition, selection would favor a similar mechanism in the other. This could lead to a polychromatic greenbeard cooperative system in a spatially structured community. The question of the general conditions that favor the evolution and stability of kin recognition is different than that which is addressed here (reviewed in (Penn and Frommen, 2010)). We simply show that there are conditions under which a recognition system could provide a fitness benefit.

The simulation model used for this research was inspired by an investigation into the spatial arrangement of microbial communities with different types of ecological interactions (Momeni et al., 2013). While the goal of our work was to compare the effects of obligate and restrictive cooperation on population growth, and not to measure spatial characteristics *per se*, it is worth noting the similarities and differences between the sets of simulated communities from the two research endeavors. Briefly, Momeni et al. described the effect of one cell type on the other as either neutral (∼), positive (↑), or negative (↑), but assumed the effect of a cell on its own type was neutral. For example: (∼,∼) was baseline cooperation in which cell types simply competed for resources; (∼, ↑) was amensalism in which one cell type harmed the other; and (↑,↑) was cooperation in which each cell type benefitted from the other.

A main conclusion from Momeni et al.’s research was that cooperation (↑,↑) led to spatial “intermixing” of lineages, while other ecological interactions led to spatial segregation. In their conceptual framework, cooperation was defined as a relationship *between* lineages and was inspired by the biological phenomenon of two auxotrophic cell types secreting products that complement the other’s needs. It therefore makes intuitive sense for the cell lineages to mix as they enhance each other’s growth. In our simulations, cooperation was defined *within* a lineage and was inspired by a different biological phenomenon, one in which a single lineage produces goods that enhance its own growth, but goods that can possibly be utilized by others, unless restricted. Comparing the models, our “obligate cooperation” (with the potential for non-cooperators to be considered cheaters) is the same as their commensal (∼,↑), except in our case, the lineage providing the growth benefit also helps its own growth. Similarly, our “two simple greenbeards” is the same as their (↑,↑) cooperation, except again, our lineages also enhance their own growth.

We did not calculate an intermixing index, but we can ask qualitatively, what kind of spatial arrangements were generated by the cooperative interactions in our simulations? Most of our simulated colonies showed strong spatial segregation. The two scenarios that showed the most intermixing between lineages were: (1) an obligate cooperator with a non-cooperator (Figure 2b), and (2) two simple greenbeards (Figure 5A); however, neither of these scenarios appeared to show as much mixing as the (↑,↑) cooperation in the Momeni et al. simulations. Notably, these are the only two cases in which none of the lineages can restrict the growth benefits of cooperation. Using their definitions, restricting cooperation simply reverts a lineage to neutral (∼) in its effect on the other strain. Thus, most of our simulations fall within the “non-cooperation” designation of their framework and have the spatial assortment associated with it. It thus appears that the amount of intermixing expected in a cooperative microbial community depends strongly on the type of cooperation: restricting the benefits of cooperation to like kinds can lead to enhanced segregation, rather than the cooperative intermixing previously reported.

While the details of the simulations and the parameter values used here may not be representative of all, or even most, microbial communities, the qualitative result, that there are possible scenarios that favor a form of “kin recognition” in spatially structured microbial communities, may be more general. They may also shed light on recent observations in yeast and bacterial species that suggest specificity in social traits may exist in natural microbial populations (Brückner et al., 2020; Butaitė et al., 2017).

## Supplementary Material

### A. Simulation Method

To model cell growth occurring in continuous time, the simulation used a discrete time Monte Carlo method with a small time step Δ*t* = 0.1 and random sequential updating. A cell was selected at random for possible division. Its growth rate *r* (either *r*_*S*_ or *r*_*F*_ depending on cell type) was calculated from Eq. (1) or (2), and its probability to grow was *P* = *r*Δ*t*. A uniform random number 0 ≤ *u* < 1 was generated, and if *u* < *P*, the cell divided and a daughter cell was placed nearby. For each time step, a random cell selection was made *T* times, where *T* was the total number of cells in the colony at the beginning of the time step.

#### Initial condition

We began the simulation with a diluted inoculum of cells into a small square at the center of the bottom layer of our cubic array (see Fig. 1). The size of the cubic array was *N*_*c*_ = 50 cells (x-direction) by 50 cells (y-direction) with a height of *N*_*z*_ = 100 cells (z-direction). The initial square had dimensions *d* × *d*, where in our simulations *d* = 15, and a fraction *f* = 0.05 of the sites were randomly selected to fill with cells (rounded down if *f d*^*2*^ is not an integer). As random sites were selected, slow cells were placed until the allotted fraction *f*_*S*_ of slow cells was reached, after which the remaining initial cells were the fast type.

#### Local densities

To determine the effect of social interactions on a cell, we must calculate the local density of each cell type near the cell of interest. The density of slow cells, *ϕ*_*S*_, was the number of slow cells in a cube of volume (2*R*_*i*_ *+* 1)^3^ centered at the focal cell, divided by the volume (2*R*_*i*_ *+* 1)^3^. Here, *R*_*i*_ is the interaction radius. The local density of fast cells, *ϕ*_*F*_, was defined similarly. If a cell was near the boundary of the cubic array, we divided by the actual volume that surrounded the cell and was part of the array. The local densities were between 0 and 1.

Calculating the local densities could be computationally expensive because of the need to loop over many nearby cells, so we reduced how often they were calculated. At the beginning of the simulation, local densities were computed for each cell. They were recomputed only when a cell was being considered for division and its neighborhood had been altered. We used a flag for each cell to track whether its local densities were current or its neighborhood had potentially been altered by movement of nearby cells.

#### Division: Finding an empty site and cell movement

If it was determined that a cell would divide, the next step was to find an empty site for the daughter cell to be placed. Similar to Momeni et al. (2013), we first checked the eight sites immediately surrounding the selected cell in the same horizontal plane, giving priority to the four sites directly adjacent rather than diagonally adjacent to the cell (see Figure S1a). If more than one equidistant empty site was available, we randomly selected one of those empty sites. The daughter cell was placed in this empty site, adjacent to the parent, and no cells moved.

If the surrounding eight sites were full, we next considered whether the dividing cell would push nearby cells aside horizontally in a planar displacement neighborhood to make room for the daughter cell. We checked for empty sites within a square neighborhood of radius *R*_*d*_ in the same horizontal plane (see Figure S1b). Keeping track of all the empty sites that were found, we calculated the Euclidean distance to each site center. In other words, we used array coordinates to identify cell center. We found the minimum distance, and if there were multiple locations at the minimum distance, we selected between them at random. The daughter cell was placed adjacent to the parent, pushing the existing cells toward the empty site.

**Figure S1.**
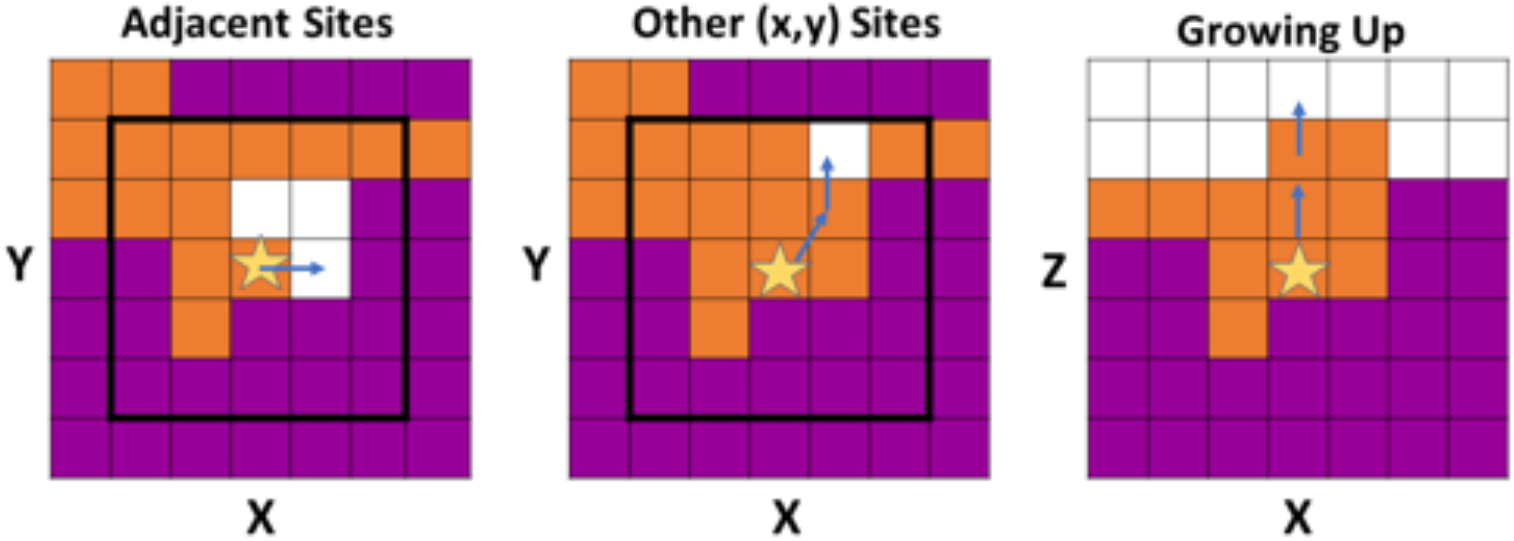
(a) A dividing cell (star) places daughter cell into a horizontally adjacent empty site. (b) A dividing cell finds the optimal path and accordingly pushes the cells on the path to the empty site, to create space to place its daughter. (c) A dividing cell places daughter directly upwards. The bold black square in (a) and (b) is the displacement neighborhood. In these figures, *R*_*d*_ = 2. This figure is adapted from Momeni et al. (2013), figure 1-S1.

The goal was to push the cells in a straight line toward the empty site, but the discreteness of the sites prevented doing this. We determined the optimal path along which to push cells as follows. From the dividing cell, we considered three adjacent cells: those in the x and y directions and the diagonally adjacent cell. Creating a vector from the dividing cell to each of those three cells, we determined which vector was most parallel to the vector from the dividing cell to the empty cell (see Figure S2). While the algorithm defined in Momeni et al. (2013) focused on distance and angle, our algorithm focused on making sure we remained as parallel as possible to the shortest path to the chosen empty site. To do this, we found the cosine of the angle between each vector and the main vector to the empty site. The cosine of each angle was calculated using the magnitudes and dot products of each of the three vectors with the main vector. The vector that created the smallest angle, or the largest cosine value, was the direction we selected to move closer to the empty site. If there was more than one direction with the maximum cosine value, we randomly chose between them. Having selected the first cell to be pushed by the daughter cell, we recorded its location. Next, we repeated the process to find the destination of the first cell being pushed toward the empty site. We continued finding cells to be pushed in the selected direction until we reached the empty site. The daughter cell was then placed next to the parent cell, and the displaced cells were pushed toward the empty site along the path that we found.

If no empty sites were found in the same plane within the displacement radius, we moved up in the z-direction (Figure S1c). Any cells above the dividing cell were pushed upward.

**Figure S2.**
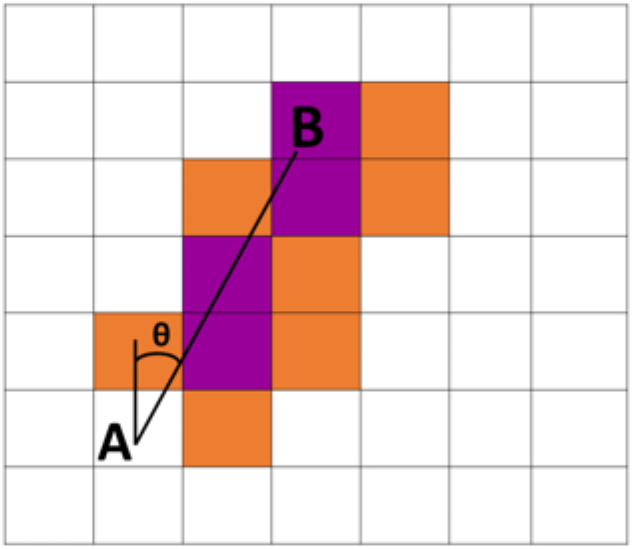
Finding the optimal path from a dividing cell at site A to an empty site B. The optimal path is along the purple sites, and the orange sites are the other neighboring sites considered.

### B. Total Proportion of Slow Cells in Various Colony Densities

Total proportion of slow cells is plotted in Figure S3. For a denser setting, we used the measurement of total proportion of slow cells at the end of the simulation, when cell totals reached 50,000 cells. This was analogous to a natural setting and is shown by the solid purple circles. We varied the maximum growth rate for slow cells as well as the initial fraction of slow cells in the inoculation droplet.

**Figure S3.**
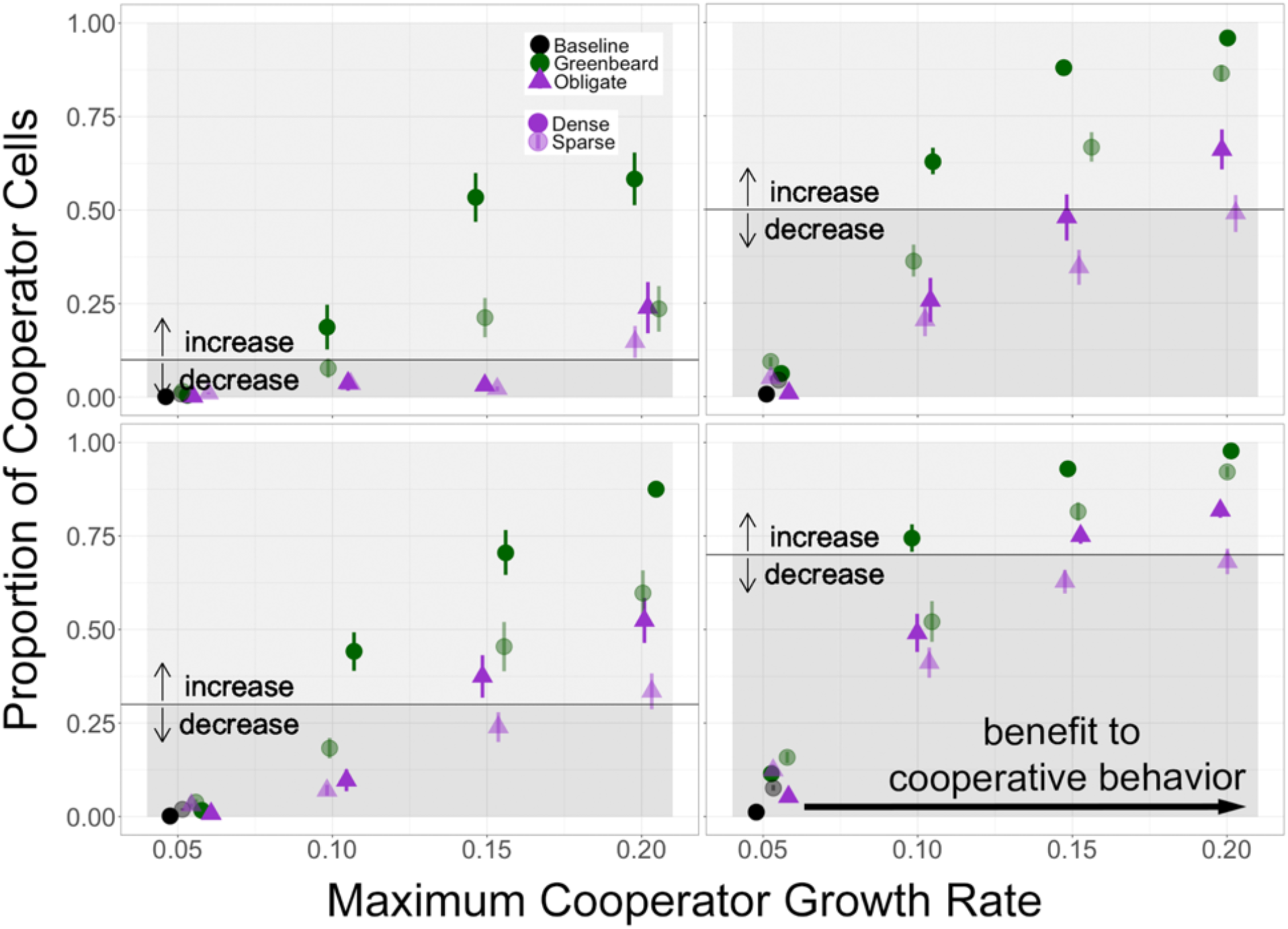
Total proportion of slow strain (S) when colonies are sparse (light purple) or dense (solid purple). Strain S are cooperators and strain F are non-cooperators; purple triangles represent obligate cooperaton and green circles represent greenbeard cooperation. The horizontal axis is the maximum growth rate for strain S. Each panel represents a different initial proportion of S. Error bars are standard error of the mean.

